# Co-immobilization of ciprofloxacin and chlorhexidine as a long-term, broad-spectrum antimicrobial dual-drug coating for polyvinyl chloride (PVC)-based endotracheal tubes

**DOI:** 10.1101/2023.04.13.534541

**Authors:** Diana Alves, Maria Olívia Pereira, Susana Patrícia Lopes

**Affiliations:** CEB - Centre of Biological Engineering, University of Minho, 4710-057 Braga, Portugal; LABBELS – Associate Laboratory, Braga/Guimarães, Portugal

**Keywords:** co-immobilization, endotracheal tube, long-term coating, polymicrobial biofilm, ventilator-associated pneumonia

## Abstract

The endotracheal tube (ETT) affords support for intubated patients, but the rising incidence of ventilator-associated pneumonia (VAP) is jeopardizing its application. ETT surfaces promote (poly)microbial colonization and biofilm formation, with a heavy burden for VAP. Devising safe, broad-spectrum antimicrobial materials to tackle ETT bioburden is needful. Herein, we immobilized ciprofloxacin (CIP) and/or chlorhexidine (CHX), through polydopamine (pDA)-based functionalization, onto polyvinyl chloride (PVC) surfaces. These surfaces were characterized and challenged with single and polymicrobial cultures of VAP-relevant bacteria (*Pseudomonas aeruginosa*; *Acinetobacter baumannii*; *Klebsiella pneumoniae*; *Staphylococcus aureus*; *Staphylococcus epidermidis*) and fungi (*Candida albicans*). The coatings imparted PVC surfaces with homogeneous morphology, varied wettability, and low roughness. Coated surfaces exhibited sustained CIP/CHX release, retaining long-term (10 days) stability. Surfaces evidencing no A549 lung cell toxicity exhibited broad-spectrum anti-biofilm activity. CIP/CHX co-immobilization resulted in better outcomes than CIP or CHX coatings, reducing bacteria up to >7 Log_10_, and modestly distressing (ca. 1 Log_10_) *C. albicans*. The anti-biofilm effectiveness of coated surfaces endured for dual biofilms, substantially preventing bacterial populations and fungi (ca. 2.7 Log_10_) in *P. aeruginosa/C. albicans* consortia. A less pronounced antifungal effect (ca. 1 Log_10_ reduction) was found in triple-species communities, but fully preventing *P. aeruginosa* and *S. aureus* populations. CIP/CHX co-immobilization holds a safe and robust broad-spectrum antimicrobial coating for PVC-ETTs, with the promise laying in reducing VAP incidence.

## 1. Introduction

The success of the current clinical practice is highly dependent on the use of medical devices or implants [1]. In this regard, the endotracheal tube (ETT) has been an elemental device in providing airway patency in patients requiring mechanical ventilation e.g., in cases like surgeries using general anesthesia, critical care situations, or a traumatically compromised airway [2]. Mechanical ventilation has long been acknowledged as a life-support practice which has picked up during the latest COVID-19 pandemic, given the unprecedented number of patients that had to be admitted to intensive care units (ICU) with severe symptoms of respiratory failure [3,4]. Once placed, the ETT becomes potentially harmful to critically ill patients undergoing long-term artificial ventilation, as it impairs natural defense mechanisms (e.g., cough reflex; upwards mucosal ciliated transport; clearance mechanism operative in the airway), and causes mechanical tissue irritation due to breathing cycles [5,6]. Furthermore, the ordinary ETTs, which are often made of flexible materials (e.g., polyvinyl chloride, PVC), are devoid of antimicrobial activity, being continuously exposed to ambient air, and providing an ideal source for microbial colonization and the development of a so-called biofilm, both early and frequent phenomena contributing to the development of ventilator-associated pneumonia (VAP). Indeed, microorganisms reach the ETT’s distal end due to contaminated oropharyngeal contents or, eventually, gastric secretion reflux [7]. Accumulation and coaggregation of common commensal bacteria further promote the recruitment of opportunistic nosocomial pathogens, developing jointly a biofilm with serious implications ultimately leading to VAP [8]. The ETT biofilm correlates with VAP pathogenesis, as microbial cells encase themselves in a self-produced matrix of extracellular polymeric substances, conferring them protection against antimicrobial treatments and the host immune system [9]. The diversity of organisms readily colonizing the ETT surface, therefore, involve drug-resistant top-priority pathogens from the ESKAPE bacterial panel such as *Pseudomonas aeruginosa*, *Acinetobacter baumannii*, *Klebsiella pneumoniae,* or *Staphylococcus aureus*, but also others that may include commensal bacteria (*Staphylococcus epidermidis*) or even fungi (*Candida albicans*) with pathogenic potential, all contributing to the polymicrobial and recalcitrant nature of ETT biofilms [10–12].

VAP is the commonest life-threatening nosocomial infection in the developed world accounting for an estimated 22% of prevalence [13], and contributing to up to 50% of all cases in ICUs [14]. Its discrepant morbidity/mortality rates (16-44%) ICU/hospital length of stay (13-26.6 days) [15–17], and incidence range (11-84%) [16,18], with ICU antibiotic prescriptions nearby 50%, are all alarming indicators of clear action needed. Given the difficulties associated with finding an effective treatment for VAP, reducing ETT bioburden through ETT surfacè modification is likely the most suitable approach to avoid VAP. Despite years of technological progress, there are still limited reports on ETT coatings. Overall, studies employ active (directly compromising surface microbial colonization), passive (contamination-resistant surface with altered composition and/or pattern), or combinatorial antimicrobial and antifouling approaches, but most limitedly addressing antimicrobial activity and/or safety-of-use [19]. In addition, studies have proven insufficient in tackling polymicrobial ETT biofilms, an underappreciated issue with a significant fate for VAP. Therefore, the development of pertinent, that is, suitable and highly efficient materials endowed with broad-spectrum antimicrobial/antibiofilm activity, biocompatibility, and cost-effectiveness features, tends to be a gold solution for the next generation of ETTs, with promise in preventing VAP.

Within the wide range of surface functionalization strategies, the bio-inspired polydopamine (pDA) coating approach [20] stands out as one that fulfills all the aforementioned requirements. In this approach, substrata are simply immersed in an alkaline solution of dopamine that then self-polymerizes and gives rise to an adhesive film with a thickness in the nanometer range. Alongside its simple processing conditions, material independence, strong reactivity for secondary functionalization, and biocompatible features have triggered its application in the immobilization of different antimicrobials to impart biomaterials’ surfaces with anti-infective properties [21,22]. Chlorhexidine (CHX) has been extensively applied in different hospital protocols for oral hygiene of intubated patients to reduce the incidence of VAP [23]. CHX immobilization has shown great potential while imparting metallic surfaces with antimicrobial features against Gram-positive bacteria in the context of orthopedic infections [21]. Impregnation of ETTs with antiseptics, including CHX, has also led to auspicious antimicrobial outcomes toward drug-resistant bacteria and fungi [24–26]. Ciprofloxacin (CIP) is a second-generation quinolone with proven broad-spectrum activity against both Gram-negative and Gram-positive bacteria. Guidelines have recommended CIP as a second antipseudomonal agent in dual-combination regimens in VAP management [27], thus holding promise to be further immobilized on ETT surfaces.

This study aimed to devise a novel and safe ETT coating supporting antimicrobial activity against a wide range of VAP-relevant species that include bacteria and fungi. For that, a pDA-based functionalization strategy was explored for the co-/immobilization of CIP and CHX on PVC surfaces, being the modified surfaces physically characterized (morphology, roughness, wettability), and inspected for their effectiveness in inhibiting the development of single-, dual- and triple-species biofilms without compromising the viability of A549 lung epithelial cells.

## 2. Materials and Methods

### 2.1 Microbial strains and culture conditions

Five model reference bacterial strains and one fungal strain were used throughout this study, namely *Pseudomonas aeruginosa* ATCC 27853, *Klebsiella pneumoniae* ATCC 11296, *Staphylococcus aureus* ATCC 25923 (all from American Type Culture Collection), *Acinetobacter baumannii* NIPH 501^T^ (kindly provided by Professor Alexandr Nemec, National Institute of Public Health in Prague) and *Staphylococcus epidermidis* CECT 4183 (from the Spanish Type Culture Collection). The fungal reference strain *Candida albicans* SC5314 was also used. All bacterial/fungal strains were stored at 80 °C ± 2 °C in a broth medium with 20% (v/v) glycerol.

All microbial strains were plated from the frozen stock solutions and first streaked on a plate containing tryptic oy agar (TSA, Liofilchem) or a Sabouraud dextrose agar (SDA, Liofilchem) plate, respectively for bacteria and fungi. After incubation (37 °C for ± 24 h), some colonies were then collected from the agar plates and grown overnight in batches of tryptic soy broth or Sabouraud dextrose broth (TSB/SDB, Liofilchem) at 37 °C under agitation (120 rotations per minute, rpm) on a horizontal shaker (Biosan OS-20). Bacterial and fungal cells were harvested by centrifugation (9000 *g*, 5 min) and washed in sterile saline solution (0.9% w/v NaCl) or phosphate-buffered saline (PBS, pH 7.4), respectively. The concentration of bacterial suspensions was then adjusted by measuring the absorbance at 620 nm (EZ Read 800 Plus, Biochrom) and using previously established standard curves, while fungal suspension concentration was adjusted after estimating cell numbers in a Neubauer counting chamber.

### 2.2 Preparation of antimicrobial stock solutions

Stock solutions of two antimicrobial agents, CIP and CHX (both from Sigma-Aldrich, St. Louis, MO, USA) were prepared by dissolving the powders in hydrochloric acid (HCl 0.2 M) or absolute ethanol and further diluted in growth medium or PBS, depending on the procedure performed (susceptibility testing and compounds release, respectively). Storage was according to the manufacturer’s instructions.

### 2.3 PVC preparation and functionalization

For surface functionalization, 1 mm-thickness unplasticized sheets of PVC (purchased from Goodfellow Cambridge Ltd, Cambridgeshire, UK) were cut into 1x1 cm-squared coupons. Before surface modification, coupons were subjected to an ultrasonic cleaning treatment in commercial detergent (Sonasol, Henkel Ibérica, Portugal) for 5 min to remove any traces of impurities and grease. After being thoroughly rinsed with distilled water, cleaned PVC surfaces were sterilized with 70% v/v ethanol for 30 min, rinsed with sterile ultrapure water, and finally irradiated with UV light for 1 h. To confer PVC surfaces with antimicrobial features, PVC functionalization with CIP and/or CHX was performed using a previously reported one-step approach for the immobilization of CHX on stainless steel [21], the so-called mussel-inspired coating strategy. The methodology scheme used for PVC functionalization is depicted in Scheme 1. Briefly, PVC coupons were firstly immersed in 7 mL of a 2 mg/mL solution of dopamine (Sigma) freshly prepared in 10 mM bicine buffer pH 8.5 (Sigma) for 18 h at room temperature under agitation (70 rpm). After dopamine polymerization, a pDA film/coating was allowed to deposit on th PVC surface (Scheme 1A). In regards to PVC functionalization with antimicrobials, 2 mg/mL dopamine was added to CHX and/or CIP (that is, alone or combined) at concentrations ranging from 0.5 to 2 mg/mL, which were dissolved together in 10 mM bicine buffer pH 8.5. The PVC coupons were simply immersed in this solution, following the abovementioned conditions. PVC surfaces modified with CIP, CHX, or CIP/CHX combined were obtained following dopamine polymerization and antimicrobial immobilization (Scheme 1B). All modified PVC surfaces were then rinsed with sterile ultrapure water and air-dried for further application. With this functionalization technique, it was expected that compounds would be incorporated throughout the full thickness of the pDA film [28]. For comparison purposes, unmodified PVC (before pDA coating or antimicrobial(s) immobilization) and pDA-coated surfaces were used in further assays.

**Scheme 1.**
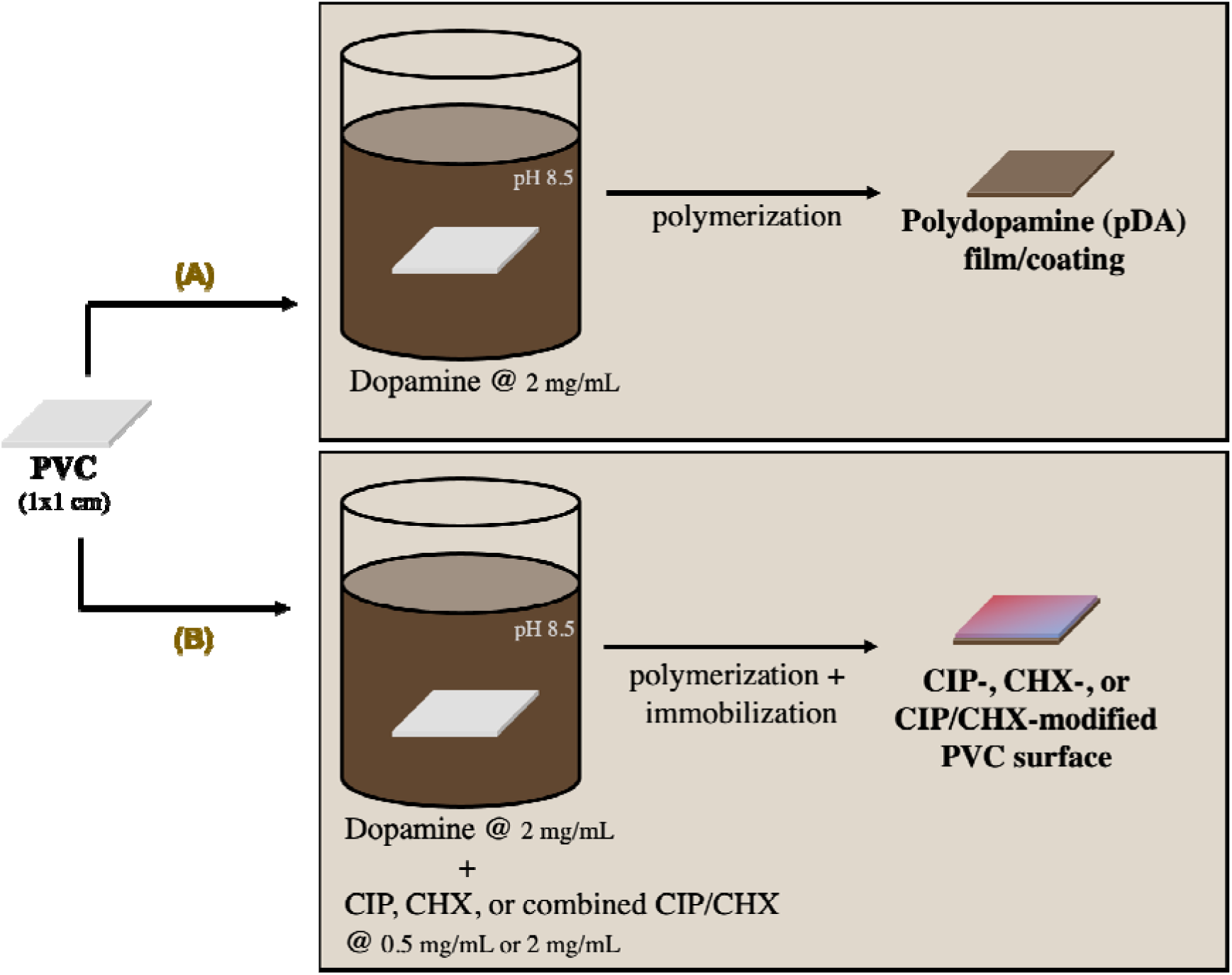
Scheme of the methodology used in the PVC functionalization with CHX and/or CIP. (A) PVC immersion in an alkaline solution (pH 8.5) of dopamine (at 2 mg/mL) resulted in its polymerization and subsequent deposition of an adhesive film called polydopamine (pDA). (B) CHX and/or CIP immobilization was performed by immersion of PVC on dopamine (at 2 mg/mL) combined with each or both compounds (at 0.5 mg/L or 2 mg/mL), resulting in CIP-, CHX-, or CIP/CHX-modified PVC surfaces upon dopamine polymerization and antimicrobial(s) immobilization.

### 2.4 Antimicrobial activity of immobilized compounds

To inspect whether CHX and/or CIP retained antimicrobial activity upon immobilization, the antimicrobial properties of the modified surfaces were performed as previously reported [21]. Briefly, each microbial suspension was adjusted to ∼1 × 10^6^ CFU/mL as described above, and 20 μL of the microbial culture was placed at the top of each unmodified/modified PVC surface and incubated at 37 ^°^C (static conditions). The suspension was allowed to dry on the top of unmodified/modified PVC surfaces, being then placed with the face exposed to microbial culture in contact with agar (TSA/SDA). Agar plates were then incubated at 37 ^°^C for 24 h, and the presence or absence of microbial growth (contact-killing) on agar was further inspected through visual observation. The antimicrobial activity of CIP- and CHX-modified PVC surfaces were also evaluated using an adaptation of the Kirby-Bauer disc diffusion method [29]. For that, modified (vs unmodified) surfaces were placed on top of TSA/SDA plates previously streaked with each microbial suspension adjusted to ∼1 × 10^8^ CFU/mL, which were then incubated (37 ^°^C) for up to 72 h. The presence or absence of an inhibition zone was further investigated and annotated.

### 2.5 Surface characterization

#### 2.5.1 Surface morphology

Surface morphology was observed by Scanning Electron Microscopy (SEM) using an ultra-high resolution Field Emission Gun SEM (FEG-SEM, NOVA 200 Nano SEM, FEI Company). Before analysis, samples were first covered with a very thin film (5 nm) of Au–Pd (80–20 wt%). Topographic images were obtained with a secondary electron detector using the following parameters: acceleration voltage of 10 kV, ∼5 mm stage distance, and 10 000 × and 50 000 × magnification.

#### 2.5.2 Surface roughness

Atomic Force Microscopy (AFM) measurements were performed at room temperature using a CSI – Nano-Observer Atomic Force Microscope, operating in tapping mode. The scanning area per sample was fixed at 10 μm × 10 μm at 512-pixel resolution. Surface morphology and roughness analysis were then conducted using Gwyddion software.

#### 2.5.3 Surface wettability

Surface wettability was investigated by measuring the static water contact angle, using a sessile drop method, in an automated contact angle measurement apparatus (OCA 15 Plus, Dataphysics, Germany) that allows image acquisition and data analysis. Contact angles were measured using 3 μL drops of ultrapure water.

### 2.6 CHX and CIP release profiles

To ascertain the release profiles of both antimicrobial compounds (CIP and CHX), the CIP- and CHX-modified PVC coupons were placed in 6-well microtiter plates (Orange Scientific, USA) to which 4 mL of PBS (pH 7.4) were added, and then incubated at 37 °C under constant agitation at 120 rpm. The PBS was all collected and refreshed at different time points (every 24 h up to 10 days). The amount of released CHX or CIP was then determined by ultraviolet-visible spectroscopy (UV–Vis), measuring the absorbance at 255 and 270 nm, respectively. The absorbance values were then converted to concentration values using calibration curves previously established.

### 2.7 Cytotoxicity determination

Cytotoxicity was evaluated on human lung epithelial A549 cells (ATCC CCL-185), according to ISO 10993-5:2006 [30]. Cells were grown in Dulbecco-modified eagle medium (DMEM, Biochrom) supplemented with 10% fetal bovine serum (FBS, Gibco) and 1% protective antibiotic solution (ZellShield™, Biochrom) at 37 °C and 5% CO_2_. Once confluence was achieved, cells were detached using trypsin-EDTA solution (0.25%/0.02%) in PBS without Ca^2+^ and Mg^2+^ (PAN Biotech GmbH, Germany), and 100 μL of cell suspension adjusted to 1 × 10^5^ cells/mL, was transferred per well to a 96-well microtiter plate. In parallel, unmodified-/modified-PVC surfaces were inserted in 24-well plates and 1 mL of supplemented DMEM was added to each well. Both plates with cells and surfaces were incubated at 37 °C and 5% CO_2_ for 24 h. After this period, the supernatant was removed and 100 μL of the medium that was in contact with the surfaces was added. Fresh-supplemented DMEM was also added as a positive control. The plate was then incubated (37 °C; 5% CO_2_) for an additional 24 h. In the dark, 20 μL of MTS (3-(4,5-dimethylthiazol-2-yl)-5-(3-carboxymethoxyphenyl)-2-(4-sulfophenyl)-2H-tetrazolium) inner salt (Promega) was added to each well and the plate was further incubated for 1 h at 37 °C, 5% CO_2_. The absorbance of the resulting solution was measured at 490 nm. The percentage of cell viability was calculated by the absorbance ratio between the cell growth in the presence of the coating and the control growth (cell growth in supplemented DMEM).

### 2.8 Anti-biofilm performance of modified PVC surfaces against single species and polymicrobial consortia

The ability of the prepared modified surfaces to prevent biofilm formation was evaluated against single-species and polymicrobial cultures encompassing the aforementioned bacterial and fungal species.

Overnight cultures were used to prepare microbial inoculum with approximately 1 × 10^6^ CFU/mL in TSB (bacterial cultures) or RPMI 1640 (fungal culture). For dual and triple-species consortia, a proportion of 1:1 and 1:1:1, respectively, of each suspended bacterial/fungal inoculum was used. Unmodified- and modified-PVC squared coupons were then inserted into a 24-well microtiter plate (Orange Scientific, USA) and inoculated with 1 mL of microbial suspensions prepared as described. As stated above, unmodified PVC and pDA-coated surfaces were used for comparison purposes. The plates containing the coupons were kept at 37 °C for 24 h, under static conditions. Coupons were then washed twice with saline solution (for bacteria) or PBS (for fungi) and transferred to new wells filled with 1 mL of saline solution or PBS. Adhered bacterial cells were removed from the PVC coupons by ultrasonic bath treatment in a Sonicor SC-52 (Sonicor Instruments) operating at 50 kHz, for 6 min, while fungal-adhered cells were removed by scrapping the surfaces (parameters previously optimized). The biofilm samples, after detached from surfaces, were afterward collected, and vortexed to disrupt possible cell aggregates, 10-fold serially diluted, and plated into appropriate agar plates that were incubated for 16-48 h at 37 °C in an aerobic incubator before biofilm cell enumeration. Estimation of bacterial and fungal cells was made on unspecific culture agar media (TSA and SDA, respectively) in single-species consortia. For microbial isolation in polymicrobial consortia, specific solid growth media were employed. *Pseudomonas* Isolation Agar (PIA, Sigma) was used to specifically select *P. aeruginosa* when in combination with *S. aureus*, *S. epidermidis*, and *K. pneumoniae,* while TSA supplemented with 10 mg/L of amphotericin B (Sigma) was used to isolate *P. aeruginosa* while suppressing *C. albicans* growth. In turn, *C. albicans* was grown in SDA supplemented with 30 mg/L of gentamicin (Nzytech) to suppress *P. aeruginosa* growth. *S. aureus* and *S. epidermidis* were both selected on Mannitol Salt Agar (MSA, Liofilchem) and *K. pneumoniae* on Klebsiella ChromoSelect Selective Agar Base (Sigma). Unlike for the other consortia, no selective media for *A. baumannii* was found, so the total number of adhered cells in TSA was compared with the growth of *P. aeruginosa* only in PIA (allowing to infer about the differential amount of *A. baumannii* cells in consortia.

### 2.9 Interpretation of anti-biofilm performance outcomes

The outcomes from the antimicrobial performance resulting from the co-immobilization of CHX and CIP were classified as synergism or facilitation according to a previously reported methodology [31]. An outcome was classified as “synergism” when the coating with combined compounds was able to prevent the adhesion of a greater fraction of microorganisms than expected if the compounds were acting independently. The term “facilitation” was used to classify an outcome in which the coating with combined compounds was better than the best of the single compounds, but not better than if the antimicrobials were acting independently. For their calculation, synergism was considered when the equation Log_10_ (S_C_) - Log_10_ (S_CHX_) - Log_10_ (S_CIP_) + Log_10_ (S_MIX_) < 0 was valid. In turn, facilitation was considered when both equations Log_10_ (S_MIX_) - Log_10_ (S_CHX_) < 0 and Log_10_ (S_MIX_) - Log_10_ (S_CIP_) < 0 were valid. In these equations, S_C_ refers to the microbial density obtained in the control (unmodified PVC surfaces), and S_CHX_, S_CIP_, and S_MIX_ to the surviving cell density after being in contact with surfaces functionalized with CIP, CHX, or both antimicrobials combined (CIP/CHX), respectively.

### 2.10 Statistical analysis

Statistical analysis and graphs were performed using the GraphPad Prism software package (GraphPad Software version 8.2.0). Means and standard deviations (SDs) were calculated for all experimental conditions tested. Statistical analysis was carried out by one-way or 2way ANOVA followed by Tukey’s multiple comparisons, and p-values < 0.05 were considered significant. At least, three independent experiments in triplicate were performed for all experiments.

## 3. Results

### 3.1 Antimicrobial screening of CIP- or CHX-modified PVC surfaces

To inspect whether CIP and CXH would retain antimicrobial activity upon immobilization, an antimicrobial screening was made, based on the ability of CIP- or CHX-modified surfaces to prevent microbial growth by contact-killing, and by antimicrobial diffusion/release on microbial-embedded agar media. The outcomes from the contact-killing and “zone inhibition” tests are depicted in Table 1.

**Table 1.** Antimicrobial activity and qualitative release of CHX or CIP from PVC-modified surfaces. A) Microbial growth after 24 h contact with CHX- or CIP-modified surfaces, where “+” is indicative of visible bacterial growth and “–” means no visible growth observed. B) Release of CHX or CIP on solid agar, qualitatively evaluated by the presence (P) or absence (A) of an inhibition zone. pDA-coated surfaces were also tested and used for comparison

|  |  | CHX-modified PVC |  | CIP-modified PVC |  |
| --- | --- | --- | --- | --- | --- |
|  | pDA-coated surface | CHX 0.5 mg/mL | CHX 2 mg/mL | CIP 0.5 mg/mL | CIP 2 mg/mL |
| *P. aeruginosa* | + | - | - | - | - |
| *A. baumannii* | + | + | - | + | - |
| *K. pneumoniae* | + | - | - | - | - |
| *S. aureus* | + | - | - | - | - |
| *S. epidermidis* | + | - | - | - | - |
| *C. albicans* | + | - | - | + | + |

B)
|  |  | CHX-modified PVC |  | CIP-modified PVC |  |
| --- | --- | --- | --- | --- | --- |
|  | pDA-coated surface | CHX 0.5 mg/mL | CHX 2 mg/mL | CIP 0.5 mg/mL | CIP 2 mg/mL |
| *P. aeruginosa* | P | A | P | P | P |
| *A. baumannii* | P | A | A | A | P |
| *K. pneumoniae* | P | P | P | P | P |
| *S. aureus* | P | P | P | P | P |
| *S. epidermidis* | P | P | P | P | P |
| *C. albicans* | P | A | A | A | A |

Results showed that all microbial species, including bacteria and fungi, were inhibited after contacting PVC surfaces immobilized with CHX at 2 mg/mL (Table 1A). When a lower concentration of CHX (0.5 mg/mL) was used, the obtained PVC surfaces lost their ability to prevent the growth of *A. baumannii* but kept antimicrobial features against all the other species. Regarding CHX diffusion on agar plates (Table 1B), inhibition zones observed for bacterial species such as *K. pneumoniae*, *S. aureus*, and *S. epidermidis* indicated CHX release from the coupons. For its higher concentration tested (2 mg /mL), however, the released CHX was still not able to prevent the growth of *A. baumannii* and *C. albicans*. Also, surfaces functionalized with a lower concentration of CHX lost their ability to inhibit the growth of *P. aeruginosa*, when compared with those immobilized with a higher amount (Table 1B). In turn, immobilization of CIP at its highest tested concentration (2 mg/mL) generated PVC surfaces with the ability to prevent microbial growth by contact with all bacterial species investigated (Table 1A). However, no antimicrobial effect was observed against the fungal strain *C. albicans*. When coming to surfaces functionalized with a lower concentration of CIP (0.5 mg/mL), these lost the ability to prevent the growth of *A. baumannii*, as compared to the previous finding. These effects were comparable to those observed when it comes to the ability of released CIP to diffuse through agar plates and inhibit bacterial growth (Table 1B).

### 3.2 Characterization of CIP and/or CIP-modified surfaces

#### 3.2.1 Surface morphology

The surface morphology of PVC surfaces modified with CIP, CHX, and even with both combined (CIP/CHX) was characterized by SEM (Figure 1).

**Figure 1.**
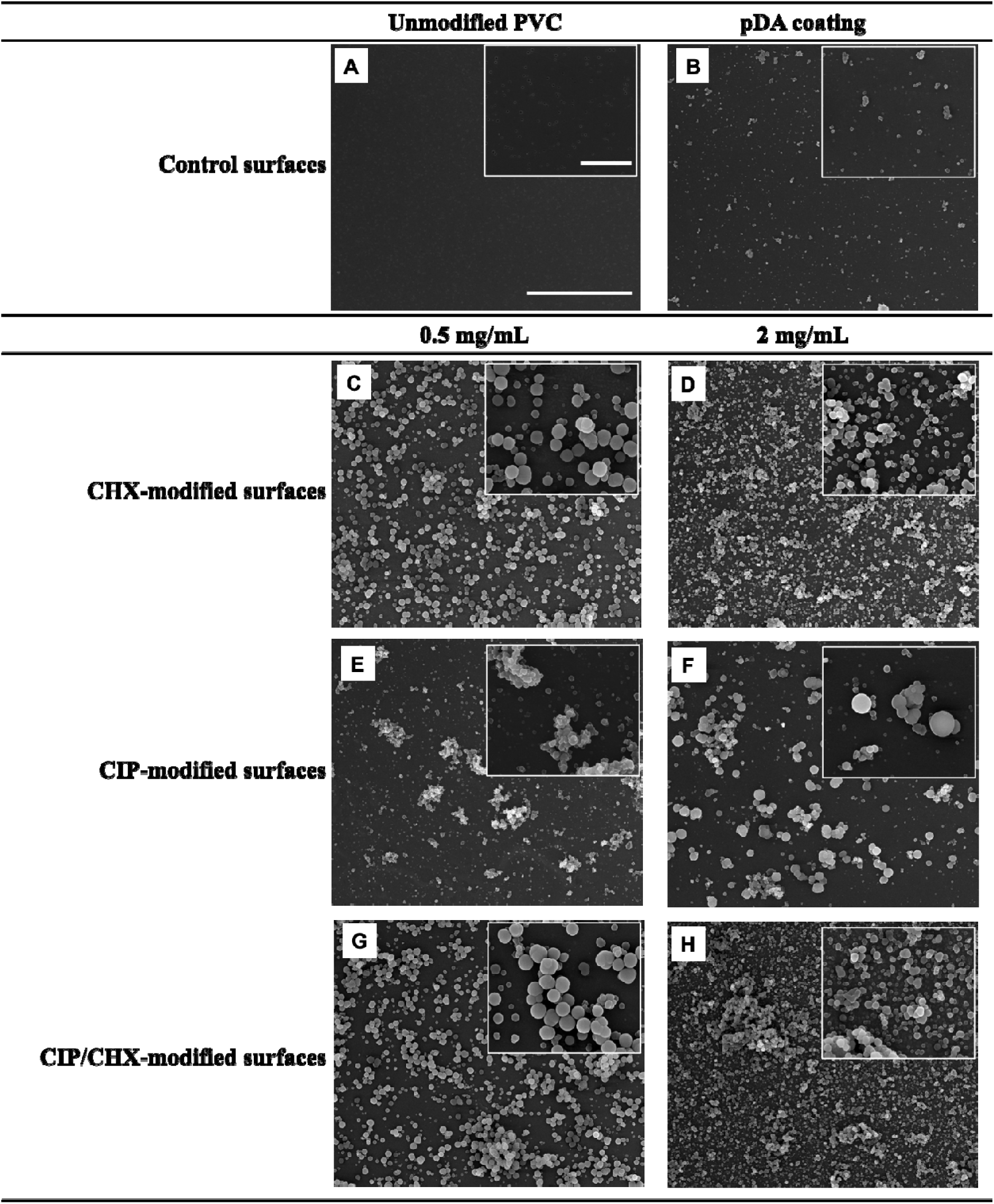
Surface morphology, obtained by SEM. (A) unmodified PVC surface; (B) pDA-coated surface; (C) PVC surface modified with CHX at 0.5 mg/mL or with; (D) CHX at 2 mg/mL; (E) CIP at 0.5 mg/mL; (F) CIP at 2 mg/mL; (G) CIP/CHX at 0.5 mg/mL, and (H) CIP/CHX at 2 mg/mL. The scale bars in the inset and image (in A) indicate 2 and 10 μm, respectively.

Bare PVC, that is, unmodified PVC surface (Figure 1A) exhibited a smooth morphology with some micropores, likely attributed to the evaporation of solvent during its production [32]. The pDA film formation on PVC surfaces (Figure 1B) has been suggested by the absence of micropores throughout the surface and the presence of self-polymerized pDA particles resulting from the bulk solution [33]. Dopamine polymerization in the presence of CHX and/or CIP resulted in altered morphologies compared to bare PVC, but preserved the structure (in clusters) of pDA-coated surfaces. In general, one-step immobilization of both compounds, either alone or combined, yielded surfaces with a more homogeneous appearance with clusters more evenly distributed throughout the surfaces (Figure 1, C to H). A more complete coverage of the surface was observed for CHX (Figure 1, C and D) when compared to surfaces functionalized with CIP alone (Figure 1, E and F). Increasing the initial concentration of the compound used for PVC functionalization apparently promoted the increase in the number of agglomerates (Figure 1, D and F vs. C and E, respectively). Co-immobilization of CHX and CIP (CIP/CHX; Figure 1, G and H) yielded surfaces with morphology and coverage similar to the ones incorporating CHX alone.

#### 3.2.2 Surface roughness and wettability

Changes in surface roughness and wettability caused by the different immobilization approaches were investigated by AFM and by measuring the static water contact angles, respectively (Figure 2).

**Figure 2.**
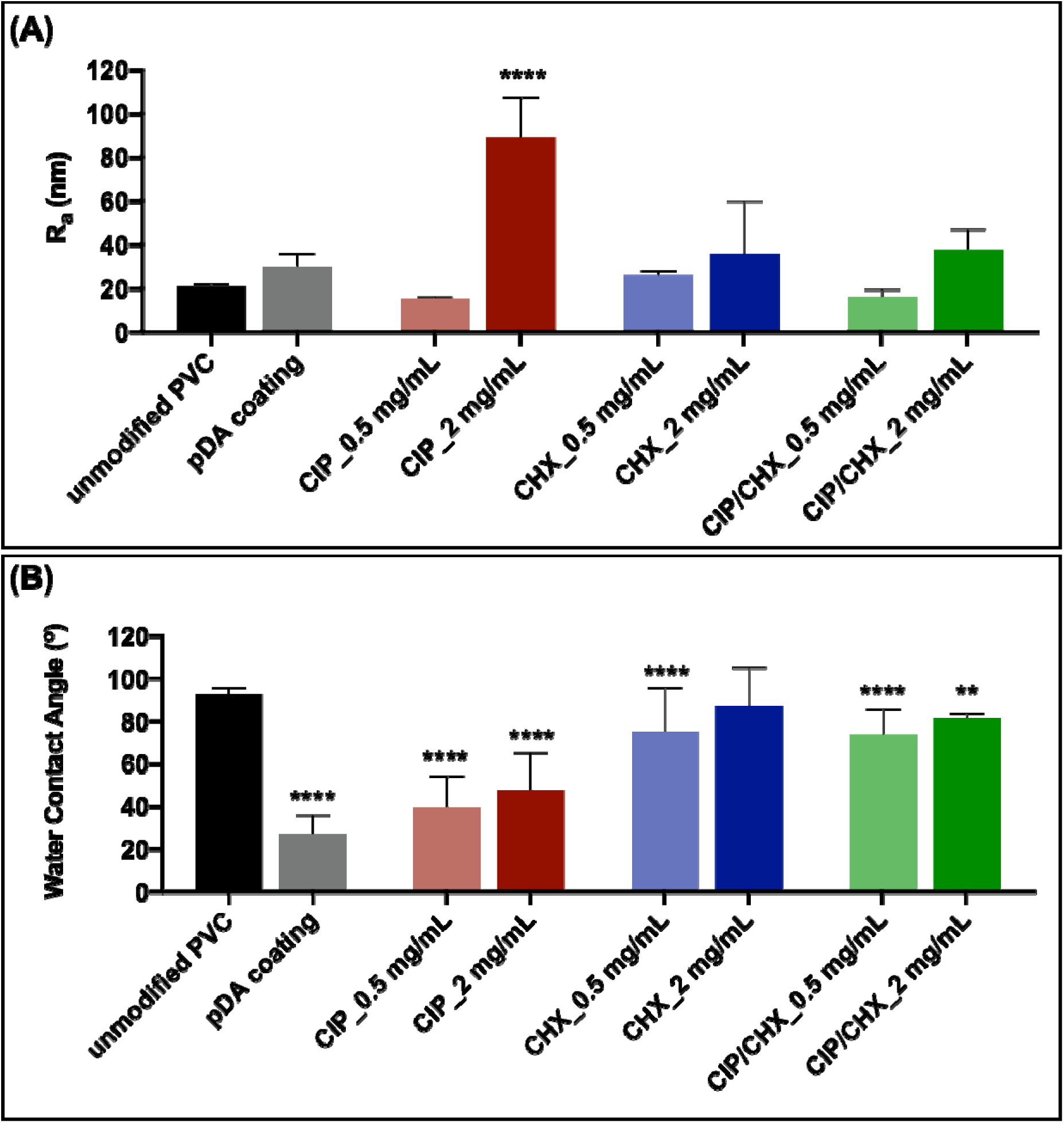
Surface roughness and wettability. (A) Average roughness and (B) static water contact angles of PVC surfaces before and after immobilization with CIP or CHX or CIP/CHX at concentrations of 0.5 and 2 mg/mL. Unmodified PVC and pDA-coated surfaces were used for comparison. The results are shown as mean ± SDs. (****) p < 0.0001, vs. unmodified PVC, one-way ANOVA, Tukey’s multiple comparison test.

The average surface roughness, expressed as R_A_ (Figure 2A), was determined from the pictures obtained from AFM analysis (Figure S1 in Supporting Information). Overall, results showed that pDA coating did not introduce significant changes in the PVC surface’s roughness as well as further functionalization with CIP and/or CHX. An exception was observed for PVC surfaces functionalized with CIP at 2 mg/mL, which displayed a significant increase (4-fold higher) in that parameter. Although not significantly different in most cases, results suggest an increase in the modified surfaces’ roughness as the concentration of immobilized compounds increases.

Water contact angles (Figure 2B) evidenced a hydrophobic character for unmodified PVC surfaces (92.6^ο^ ± 3.2^ο^). Dopamine polymerization imparted the PVC surfaces with hydrophilic properties, as evidenced by the significant decrease of the water contact angle to below 90^ο^ (27.3^ο^ ± 8.5^ο^), a well-established observation found on other materials functionalized with pDA [22,34]. Concerning modified surfaces, the incorporation of CHX during dopamine polymerization resulted in an increase in the water contact angle (75.3^ο^ ± 20.4° and 87.1° ± 18.0°), turning surfaces with more hydrophobic features. On the other hand, PVC functionalization with CIP did not greatly interfere with the hydrophilic properties imparted by pDA alone. Co-immobilization of CHX and CIP (CIP/CHX) yielded surfaces with wettability similar to the CHX-immobilized ones. Overall, increasing the initial concentration of the immobilized compounds, when alone or combined, slightly increased the hydrophobic properties of the modified surfaces.

### 3.3 CHX and CIP release profiles

The released amount of CHX and CIP from functionalized PVC surfaces was monitored each day for up to 10 days, by measuring the absorbance by UV-Vis and further determining the released amount of each compound through previously established calibration curves. The profiles obtained for CIP and CHX cumulative released mass are presented in Figure 3.

**Figure 3.**
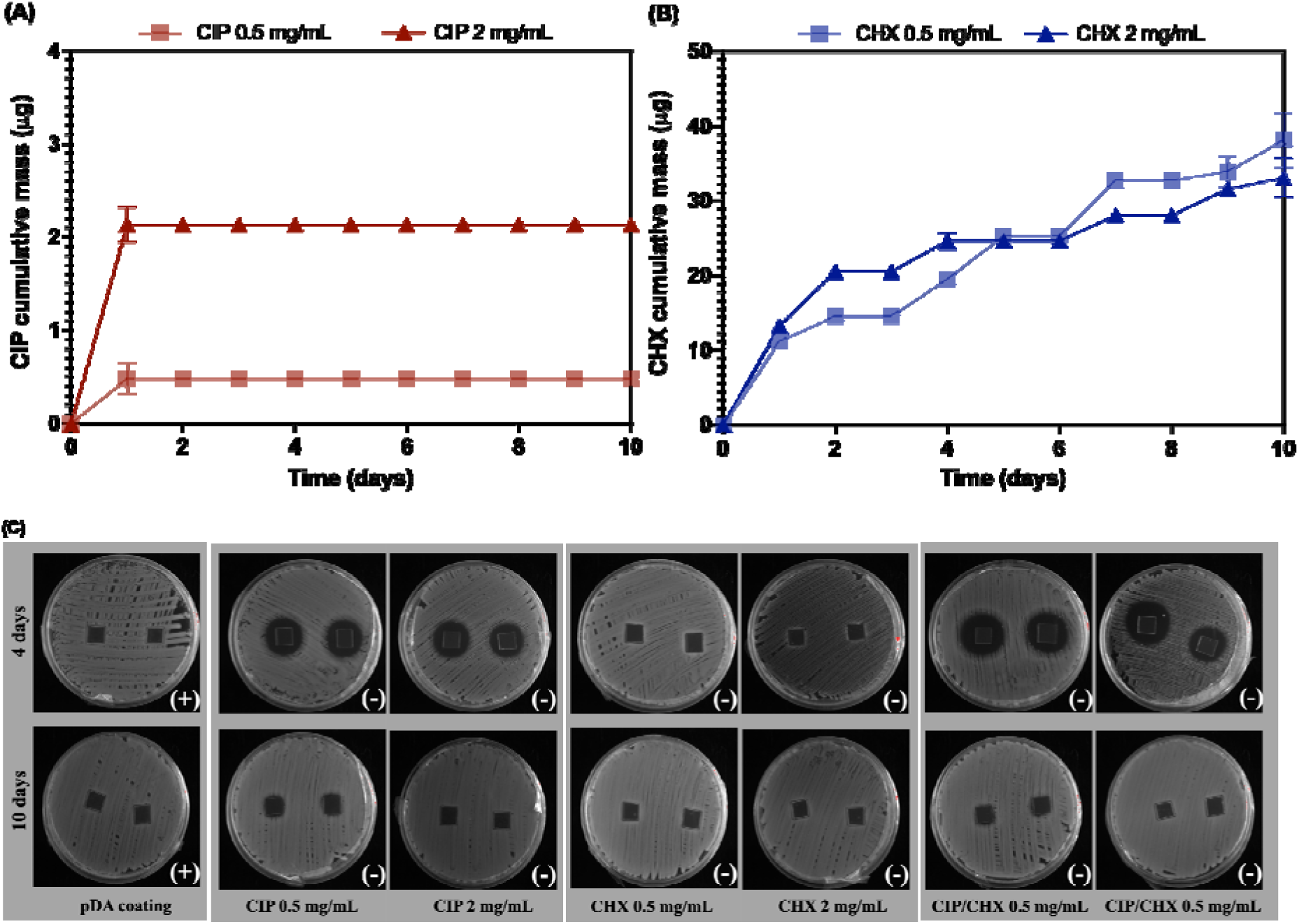
Antimicrobials release from coated surfaces. (A) Cumulative release of CIP; and (B) of CHX from PVC surfaces functionalized with each compound at 0.5 and 2 mg/mL. Panel C depicts the outcomes from contact-killing and release of CIP or CHX immobilized alone, and CIP/CHX co-immobilized on PVC surfaces at 0.5 and 2 mg/mL, at days 4 and 10 against *K*.

*pneumoniae*. Bacterial growth after 24 h contact with modified surfaces is indicated at the bottom right of each image, where “+” is indicative of visible bacterial growth and “–” means no visible growth observed. The release of antimicrobials on solid agar was qualitatively evaluated by the presence or absence of an inhibition zone. pDA-coated surfaces were also tested and used for comparison.

Overall, there was an initial burst release of CIP immobilized at both concentrations, which was observed in the first 24 h after its immobilization on PVC surfaces (Figure 3A). During the following days, CIP could not be detected (no CIP mass added). A comparable trend was noted for PVC surfaces functionalized with a higher CIP concentration (2 mg/mL), however, showing an enhanced released amount of CIP in comparison to those surfaces immobilized with a lower CIP concentration. A significantly higher cumulative mass was found for CHX (Figure 3B), as compared to CIP, with cumulative mass estimations nearby 40 µg after 10 days of release. As for CIP, a rampant rise in CHX amount was found for the early 24 h, followed by a slower release of the antimicrobial in the remaining days. Initially, the amount of liberated CHX increased as the immobilized CHX concentration also increased (< 4 h). From then on, PVC surfaces functionalized with a lower CHX concentration (0.5 mg/mL) started to release higher CHX amounts than those immobilized with CHX at 2 mg/mL.

As stated above, the apparent CIP depletion in the PBS supernatant for the period after 24 h of release, was likely attributed to a low-limit detection. To support this, the presence of released CIP, alone and combined with CHX, was further investigated on days 4 and 10, based on direct contact-killing and release approaches, as demonstrated in panel C of Figure 3. *K. pneumoniae*, shown as the most susceptible species to both CIP and CHX in broth microdilution testing (Table S1 of Supporting Information), was used so that lower antimicrobial concentrations could be detected. At day 4, all CIP-and CIP/CHX-modified surfaces exhibited contact-killing and formed inhibition zones against *K. pneumoniae*. Such results still evidenced immobilization of CIP, either alone or combined, on the PVC surfaces but in small quantities likely not detected by UV spectroscopy. The presence of CHX in the supernatant was also appraised by means of contact-killing and release assets. As expected, the absence of an inhibition zone may be a reflection of its low release concentration on day 4 (Figure 3B), which was found below its minimum microbiocidal concentration (MMC; Table S1). After 10 days of release, no inhibition zone was observed for any modified surface, but all CIP, CHX, and CIP/CHX-modified surfaces retained their contact-killing activity (as indicated by the symbol “(-)” at the bottom right of the images), evidence of an effective antimicrobial immobilization.

### 3.4 Cytotoxicity against A549 lung epithelial cells

Aiming at evaluating the toxicity of PVC-modified surfaces to lung epithelial cells, PVC surface immobilized with CIP and CHX, alone or combined, were placed in contact with a DMEM cultur medium, which was further exposed to A549 cells. The cell viability (expressed as a percentage), of indirectly contacting PVC surfaces immobilized with CIP and/or CHX is presented in Figure 4.

**Figure 4.**
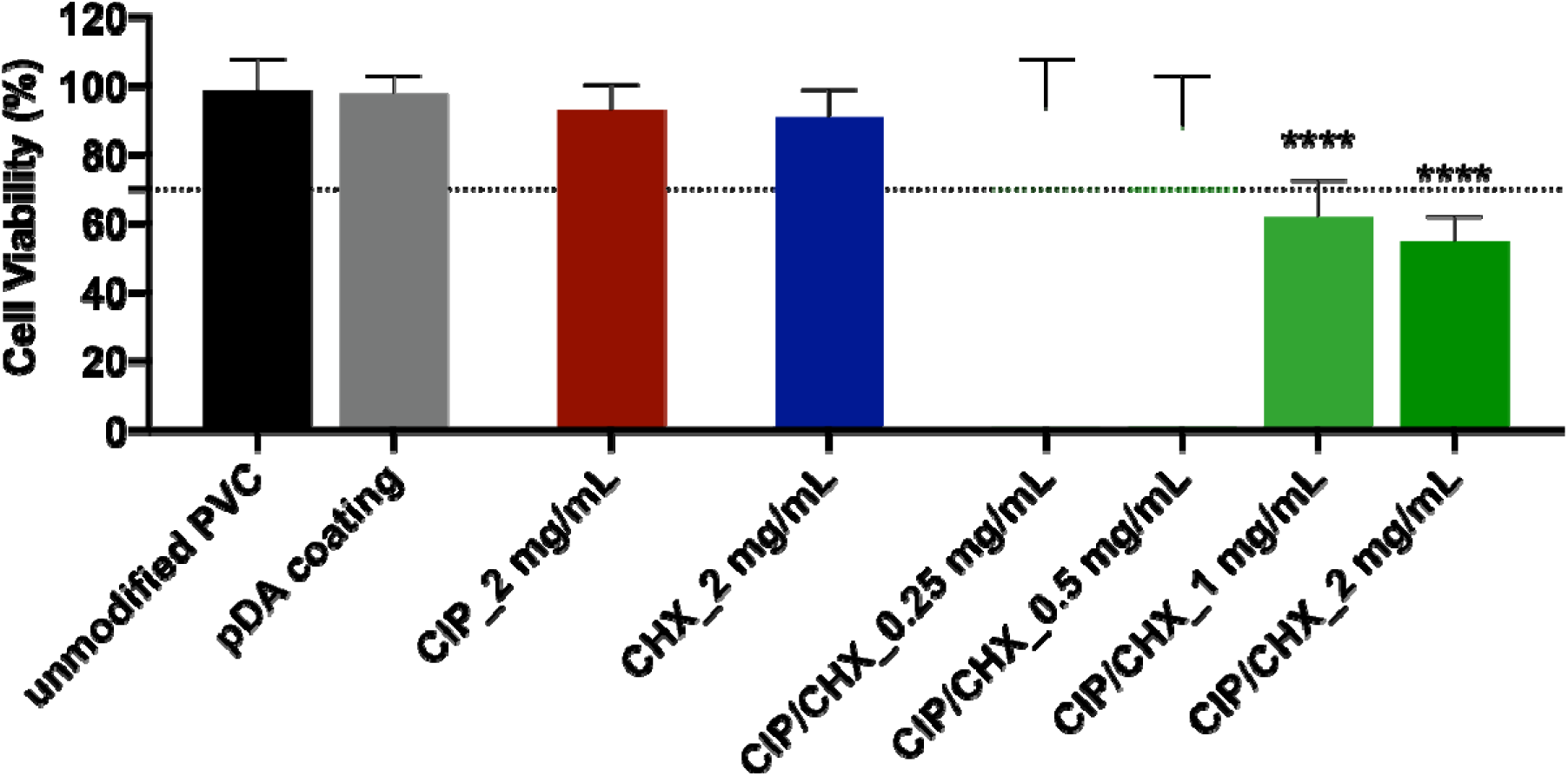
Cell cytotoxicity. Viability of A549 lung epithelial cells after indirect contact with PVC surfaces functionalized with CIP or CHX at the highest concentration tested (2 mg/mL) and with CIP/CHX at 2 mg/mL and lower concentrations (1 mg/mL; 0.5 mg/mL, and 0.25 mg/mL). Unmodified PVC and pDA-coated surfaces were used for comparison. A threshold for cell toxicity of 70% viability was used, which is indicated by a dotted line. The results are shown as mean ± SDs. (****) p < 0.0001, vs. unmodified PVC, one-way ANOVA, Tukey’s multiple comparison test.

A large percentage of A549 viable cells was obtained when in indirect contact with unmodified PVC before and after pDA coating (Figure 4). Further functionalization of PVC surfaces with CIP or CHX at 2 mg/mL also produced no significant cytotoxic effect on A549 eukaryotic cells, as compared with unmodified surfaces. In turn, CIP/CHX co-immobilization with the highest antimicrobial concentration tested (2 mg/mL) on a PVC surface resulted in a decline in the number of viable cells higher than 30%, evidence of toxicity (< 70% cell viability) toward eukaryotic cells. In a way to reduce cytotoxicity, doses lower than 2 mg/mL were tested for co-immobilization, and the viability of A549 cells exposed to DMEM medium which was put before in contact with these modified surfaces was also appraised. Cytotoxicity was found to be dose-dependent, with the highest antimicrobials concentrations (2 mg/mL and 1 mg/mL) causing toxic effects on A549 cells, contrariwise to CIP/CHX doses lesser than 0.5 mg/L, which did not affect cell viability. Based on these findings, only modified PVC surfaces with proven biocompatibility proceeded to the next anti-biofilm assays.

### 3.5 Efficacy of modified PVC surfaces in preventing single-species biofilms

As microbial adhesion is the preceding step of biofilm formation, which represents an early and recurrent event in intubated patients [8,35–37], the next stage was to appraise the effectiveness of the modified-PVC surfaces in inhibiting microbial adhesion, and so preventing the biofilm development by microorganisms commonly associated with VAP, namely *P. aeruginosa*, *A. baumannii*, *K. pneumoniae*, *S. aureus*, *S. epidermidis*, and *C. albicans*. Thus, PVC surfaces modified with CIP and/or CHX at the highest concentrations displaying no cytotoxicity (0.5 mg/mL and 2 mg/mL), were challenged for 24 h with microbial suspensions of each aforementioned species, and the number of cultivable cells that adhered to the surfaces was enumerated after detachment (Figure 5).

**Figure 5.**
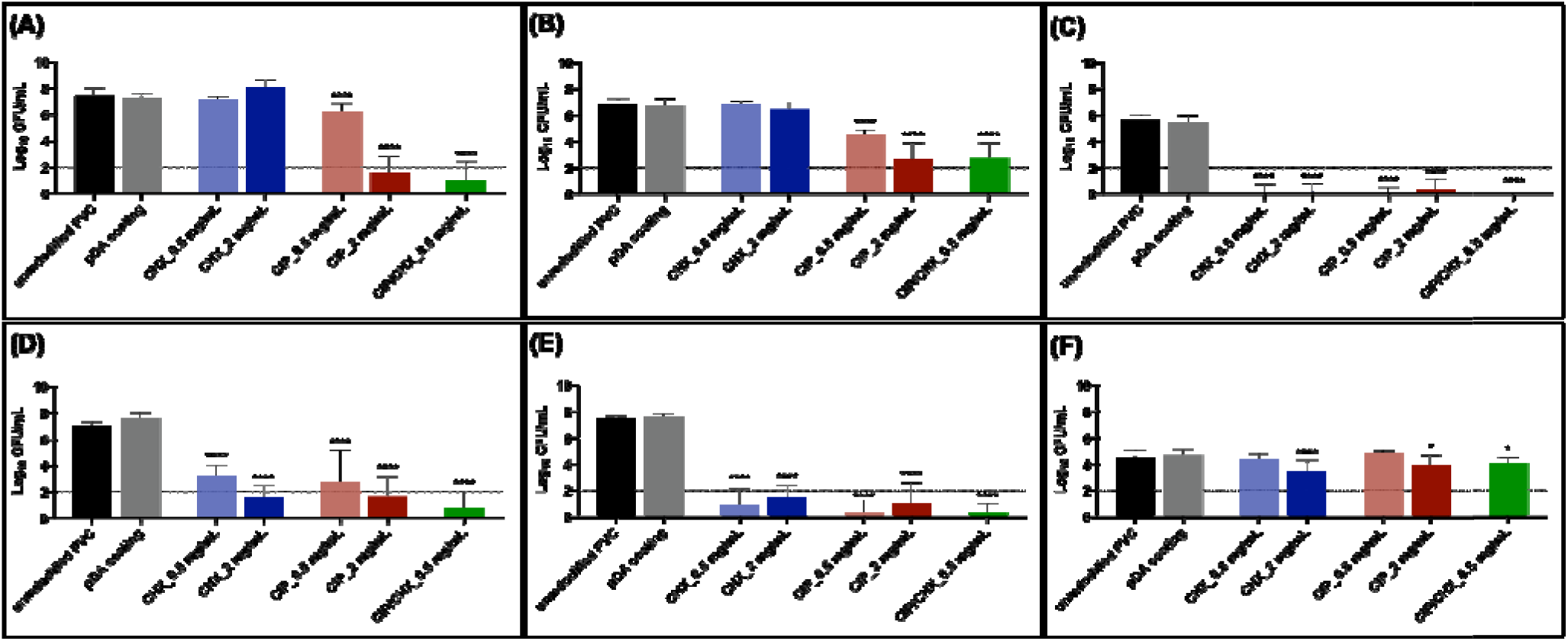
Efficacy against single-species biofilms. PVC surfaces modified with CIP and/or CHX challenged with single-species biofilms formed by *P. aeruginosa* (A), *A. baumannii* (B), *K. pneumoniae* (C), *S. aureus* (D), *S. epidermidis* (E), and *C. albicans* (F). Unmodified PVC and pDA-coated surfaces were used for comparison. The threshold value marked by the dotted lin represents the detection limit of CFU counting. The results are shown as mean ± SDs. (****) p < 0.0001, vs. unmodified PVC, one-way ANOVA, Tukey’s multiple comparison test.

Results show that the ability to form biofilms on bare/unmodified PVC surfaces relied on the microbial species, with most biofilms achieving high cell densities varying from 6.83 and 7.52 Log_10_ CFU/mL after 24 h. Lower cell concentrations, however, were found for *K. pneumoniae* (Figure 5C) and *C. albicans* (Figure 5F), reaching 5.64 and 4.57 Log_10_ CFU/mL on average, respectively. As anticipated, the pDA film on PVC had no effect on the adhesion of any microbial species, as evidenced by the similar number of adhered cells found compared to the unmodified PVC control.

Functionalization with CHX and/or CIP rendered the PVC surfaces with wide-ranging anti-biofilm activities. CHX-functionalized surfaces were not enabled to prevent biofilm formation by *P. aeruginosa* (Figure 5A) and *A. baumannii* (Figure 5B), regardless of the CHX initial concentration applied. However, significant reductions were found for the remaining biofilm populations (Figure 5, C-F). A dose-dependent reduction was observed for *S. aureus*, by 3.8 Log_10_ (CHX 0.5 mg/mL) and 5.5 Log_10_ (CHX 2 mg/mL) (Figure 5D), but also for *C. albicans* biofilms, which was significantly inhibited (∼ 1.1 log_10_) when CHX 2 mg/mL was immobilized on PVC surfaces (Figure 5F).

As for CHX, great reductions and, in most cases, a dose-dependent anti-biofilm effect were observed for CIP-functionalized PVC surfaces. For instance, *P. aeruginosa* decreased from 6.26 to 1.60 Log_10_ CFU/mL (Figure 5A), *A. baumannii* decreased from 4.60 to 2.68 Log_10_ CFU/mL and *S. aureus* from 2.73 to 1.65 Log_10_ CFU/mL (Figure 5D) when grown on PVC immobilized with CIP at 2 mg/mL compared to CIP at 0.5 mg/L. CIP immobilization at 2 mg/mL still caused a slight reduction (< 1 log_10_) in the number of fungal biofilm cells (Figure 5F). The highest inhibitory outcomes were found for biofilms of *K. pneumoniae* (Figure 5C), *S. aureus* (Figure 5D), and *S. epidermidis* (Figure 5E), with the modified surfaces, substantially impairing their developments (cell numbers were found below the thresholds for cultivability).

Interestingly, co-immobilization of CIP and CHX, exhibited a significant anti-biofilm effect against all consortia investigated. The estimated outcomes of PVC co-immobilization regarding its anti-biofilm performance (Table S2 of Supporting Information) demonstrated a facilitative effect against all microbial species and even a synergic outcome against *P. aeruginosa*, *A. baumannii*, and *C. albicans*. For better understanding, calculations showed better anti-biofilm performance of CIP-CHX co-immobilization than the best of the single compounds acting on all consortia, also enabling to prevent the adhesion of a greater fraction of microbial cells in *P. aeruginosa*, *A. baumannii*, and *C. albicans* biofilm communities than expected if the compounds were acting independently.

### 3.6 Efficacy of modified PVC surfaces against dual-species biofilms

Considering the great prevalence of *P. aeruginosa* on VAP [11,38,39], the PVC surfaces immobilized with CIP and CHX, alone or combined, were then evaluated against the biofilm formation of this species combined with each one of the other tested microbial species (Figure 6).

**Figure 6.**
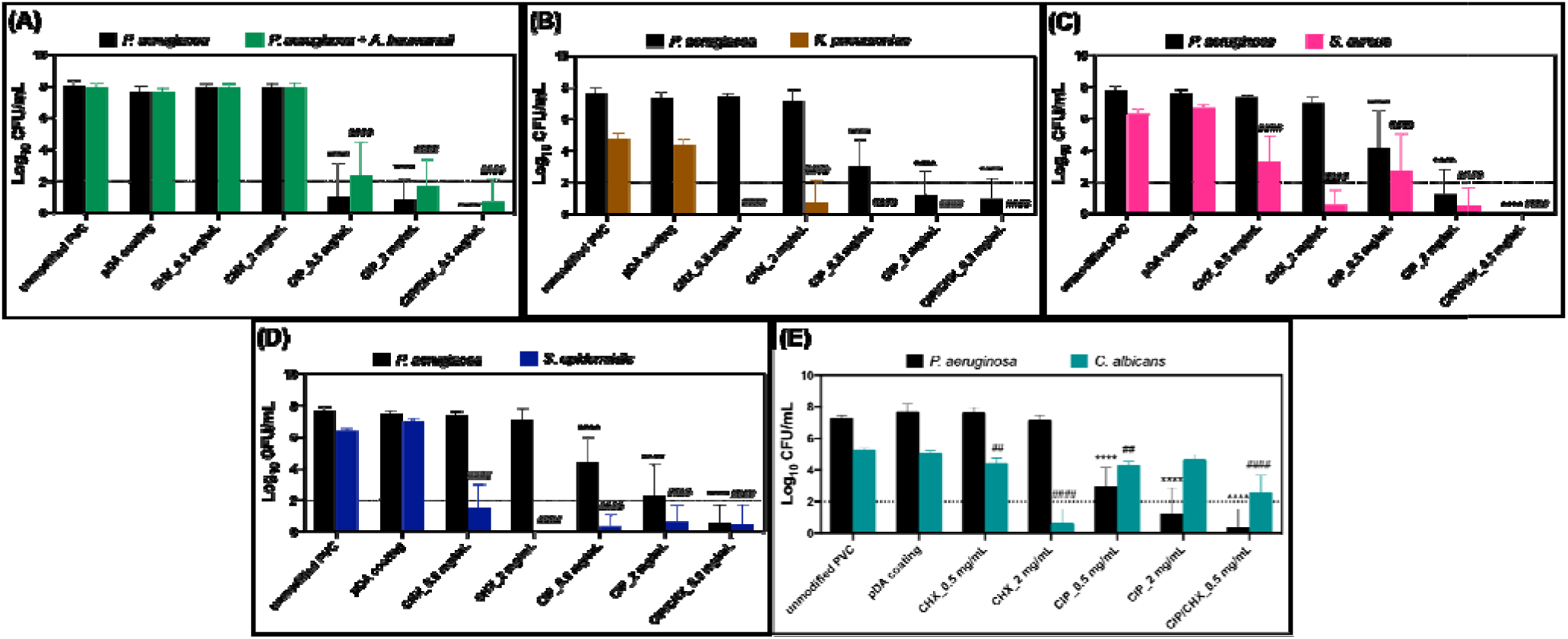
Efficacy against dual-species biofilms. Effect of PVC surfaces before and after pDA-based coating and immobilization of CIP and/or CHX against dual-species biofilms of *P. aeruginosa* and *A. baumannii* (A), *P. aeruginosa* and *K. pneumoniae* (B), *P. aeruginosa* and *S. aureus* (C), *P. aeruginosa* and *S. epidermidis* (D), and *P. aeruginosa* and *C. albicans* (E). Unmodified PVC and pDA-coated surfaces were used for comparison. The threshold value marked by the dotted line represents the detection limit of CFU counting. The results are shown as mean ± SDs. (****) p < 0.0001 vs. *P. aeruginosa* growth on unmodified PVC; and (^##^) p < 0.01 or (^####^) p < 0.0001 vs. other than *P. aeruginosa* species growth on unmodified PVC, 2way ANOVA, Tukey’s multiple comparison test.

Overall, dual-species biofilms were mostly governed by *P. aeruginosa*, regardless the PVC material where the consortia were formed. Biofilms developed on unmodified surfaces (PVC and pDA) presented cell numbers in orders of magnitude similar to those observed for *P. aeruginosa* alone (see Figure 5A), denoting low influence of the other microbial species in consortia in *P. aeruginosa* growth.

CHX-modified surfaces were able to significantly disturb all other than *P. aeruginosa* populations in the consortia (Figure 6, A-E), lowering CFU counts to below the detection limit or even leading to full growth prevention of some bacterial species, namely *K. pneumoniae* (Figure 6B) and *S. epidermidis* (Figure 6D). The absence of a specific medium for the isolation *A. baumannii* is a limitation of this study, preventing us to infer the influence of PVC surfaces on *A. baumannii* growth (when in consortia with *P. aeruginosa*). For PVC, pDA, and the coatings with CHX alone, no significant differences were detected between the number of adhered cells found on TSA or on the selective *P. aeruginosa* medium, PIA (Figure 6A), which is indicative that *P. aeruginosa* is still likely the most dominant species in these conditions (as ascertained by Figure S2, in Supporting Information). It would be expected, however, a lack of effectiveness of CHX-modified surfaces against *A. baumannii* population similar to the one observed in its single-species biofilms.

When coming to CIP-modified surfaces, drastic dose-dependent reductions were estimated for the biofilm populations, as compared with those developed on unmodified PVC surfaces. Such reductions were more pronounced for the bacterial populations in all consortia (Figure 6, A-D), as compared with the fungal population (Figure 6E). Concerning *P. aeruginosa* and *A. baumannii* consortia (Figure 6A), equitable CFU numbers were found for each bacterial population within the biofilms grown in CIP-modified surfaces, with PIA estimating about 1.0 Log_10_ and 0.9 Log_10_ CFU/mL of *P. aeruginosa* cells, whereas *A. baumannii* presented ∼1.3 Log_10_ and 0.7 Log10 CFU/mL, given by the difference between TSA and PIA counts, respectively (CIP 0.5 mg/mL vs. 2 mg/mL). As compared to their single-species biofilms (see Figure 5, A and B), the extent of adhesion of both populations in dual-species consortia was notoriously affected by CIP-modified surfaces, reducing *P. aeruginosa* growth by approximately 5.3 Log_10_ and 0.8 Log_10_ and *A. baumannii* by 3.3 Log_10_ and 2.0 Log_10_ (CIP 0.5 mg/mL vs. 2 mg/mL). When CIP was immobilized and combined with CHX, they also led to greater reductions in *P. aeruginosa* (∼1 Log_10_) and *A. baumannii* (∼2 Log_10_) populations, compared to its effect in single-species consortia.

Biofilm formation by *K. pneumoniae* and *S. aureus* on unmodified PVC and pDA surfaces was compromised by the presence of *P. aeruginosa*, as evidenced by ca. 1 log_10_ reduction in the number of biofilm cells compared to their single-species biofilm (Figure 6, B and C vs Figure 5, C and D, respectively). Further PVC functionalization with CIP and CHX, alone or combined, led to substantial decreases in biofilm cell numbers for both species (Figure 6, B and C), even fully preventing *K. pneumoniae* biofilm population, a result comparable to that observed for its single-species biofilm (Figure 6B vs Figure 5C). Similar to other bacterial species, *S. epidermidis* biofilm formation on untreated surfaces was impaired by the presence of *P. aeruginosa*, as evidenced by a reduction of approximately 1 log_10_ and 0.7 log_10_ for unmodified PVC and pDA surfaces, respectively. However, it should be taken into account that some of this reduction could be associated with its growth in the selective media (Figure S2 in Supporting Information). All the CIP- and/or CHX-modified surfaces caused a significant reduction in the number of *S. epidermidis* cells in the dual-species consortia (Figure 6D), an antimicrobial performance similar to the one obtained for its single-species adhesion (see Figure 5E).

Contrariwise to the consortia involving only bacterial species, biofilms involving *P. aeruginosa* and *C. albicans* were not always dominated by *P. aeruginosa* (Figure 6E). Despite the predominance of the bacterial population when growing on both unmodified PVC and pDA-coated surfaces, an increase (∼ 0.7 Log_10_ and 0.3 Log_10_ compared to PVC and pDA-coated surfaces, respectively) in the fungal ability to adhere to CIP-modified surfaces were observed relatively to *P. aeruginosa*, which endured in CIP/CHX co-immobilized surfaces. Despite this differential growth among both bacterial and fungal populations in the consortia, co-immobilization of CIP and CHX in PVC surfaces also caused a significant reduction of ca. 2.7 Log_10_ on the number of adhered fungi, which was quite near the limit threshold for CFU counting. The antimicrobial performance of this coating strategy against *P. aeruginosa* in this consortium was similar to or better than the one obtained for its single-species adhesion. A major reduction was also found for bacterial/fungal biofilms grown on PVC functionalized with CHX at the highest concentration (2 mg/mL), which caused a 4.7 Log_10_ reduction in fungal cells (as opposed to the 1.1 Log_10_ reduction achieved for its single-species adhesion).

Co-immobilization of CIP and CHX proved to result in a synergic effect against *P. aeruginosa* for all consortia investigated, according to the calculations in Table S3 of Supporting Information. A similar synergic effect endured against *C. albicans* and a facilitative one against all other microbial species in consortium with *P. aeruginosa*.

### 3.7 Efficacy of modified PVC surfaces against triple-species biofilms

To better mimic the true polymicrobial (that often includes inter-kingdom) nature of VAP infection and evaluate the dual-drug CIP/CHX coating strategy’s successfulness in a more complex (and eventually, worsening) scenario, its effectiveness in preventing the development of a triple-species biofilm was investigated. Biofilms comprised of three populations involving Gram-negative (*P. aeruginosa*) and Gram-positive (*S. aureus*) bacterium, but also a fungal species (*C. albicans*) were grown on CIP/CHX-modified surfaces, and their CFU number estimated after 24 h (Figure 7).

**Figure 7.**
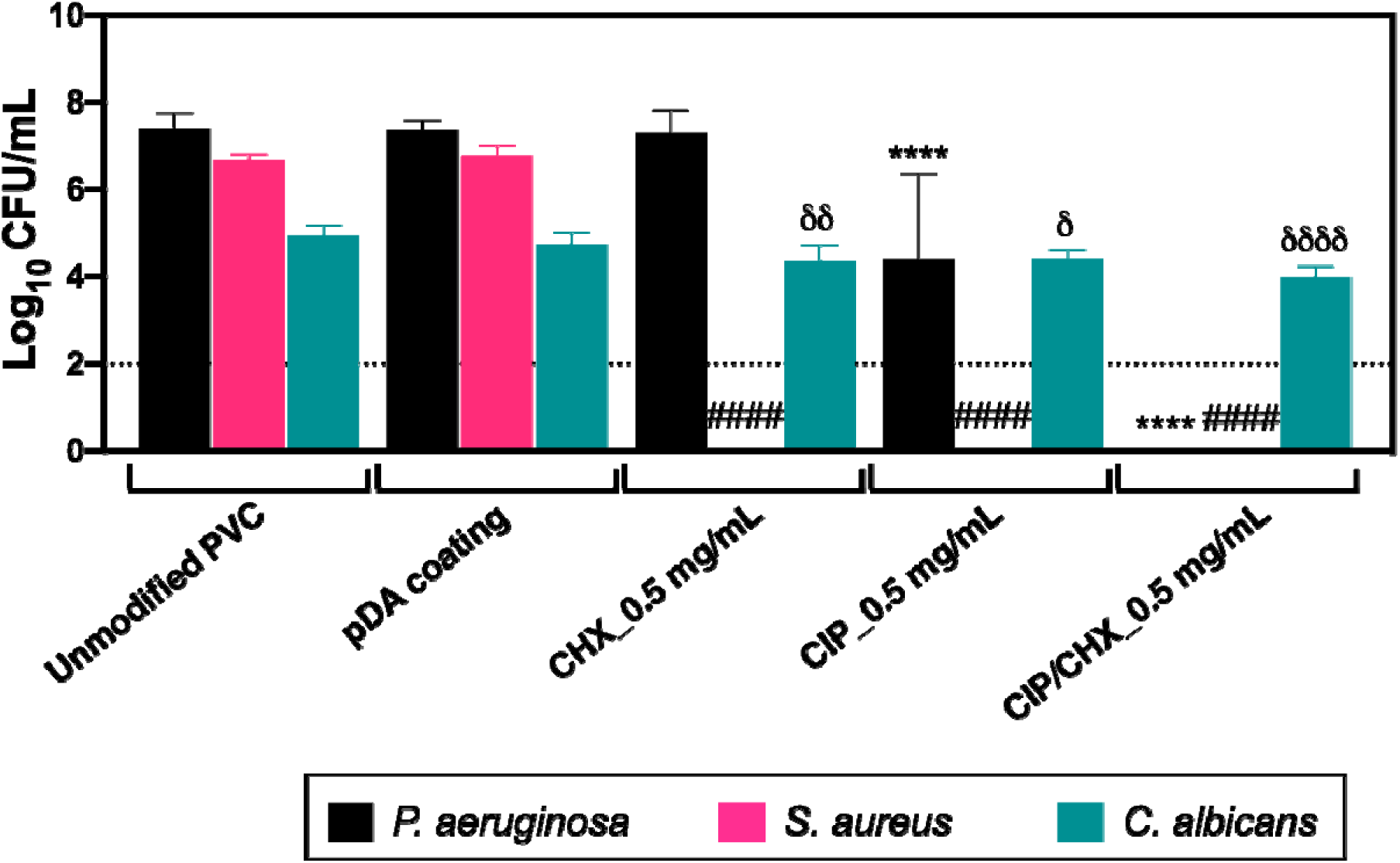
Efficacy against triple-species biofilms. Effect of PVC surfaces before and after pDA-based coating and immobilization of CIP and/or CHX against triple-species biofilms of *P. aeruginosa, S. aureus*, and *C. albicans*. Unmodified PVC and pDA-coated surfaces were used for comparison. The threshold value marked by the dotted line represents the detection limit of CFU counting. The results are shown as mean ± SDs. (****) p < 0.0001 vs. *P. aeruginosa* growth on unmodified PVC; (^####^) p < 0.0001 vs. *S. aureus* species growth on unmodified PVC; and (^∂^) p <0.05, (^∂∂^) p < 0.01 and (^∂∂∂∂^) p < 0.0001 vs. *C. albicans* species growth on unmodified PVC 2way ANOVA, Tukey’s multiple comparison test.

The ability of *P. aeruginosa* or *C. albicans*, grown in triple consortia, to adhere to control unmodified PVC and pDA-coated surfaces, was not compromised by the presence of other microbial species in the biofilm, as compared to their single-species adhesion (see Figure 5, A and F). On the other hand, minor decreases (< 1 Log_10_) occurred for *S. aureus* populations (Figure 7 vs Figure 5D), in a similar way to when it was combined with *P. aeruginosa* in dual-species consortia (see Figure 6D). All modified surfaces, immobilized with CIP and CHX alone or combined, contributed to significantly impair the adhesion of *S. aureus*. In turn, PVC functionalized with CHX alone had no effect against *P. aeruginosa* population, as expected, but its growth was considerably disturbed when biofilms were developed on CIP- and CIP/CHX-modified surfaces, even resulting in pseudomonal prevention for CIP/CHX-modified surfaces (Figure 7). However, the anti-biofilm effect of all coatings was not demonstrated for *C. albicans* populations, as attained for bacterial populations in the consortia. Furthermore, the modified surfaces had a more modest antifungal effect against triple-species biofilms when compared to that attained in dual-species *P. aeruginosa/C. albicans* biofilms (Figure 6E). Even so, the dual-drug CIP/CHX coating was promising in substantially affecting bacterial populations inside the consortia, and even modestly reducing *C. albicans* cell numbers by about 1 Log_10_ (compared to unmodified PVC).

## 4. Discussion

Every time an ETT is used for invasive mechanical ventilation, there is a raise of about 6 to 20 folds in the risk of ventilated patients acquiring VAP [40,41]. This increased risk has been substantially related to microbial proliferation on the ETT surface that rapidly evolves for biofilm formation and eventually leads to complete occlusion of the ETT and spreading of pathogens to the lower airways, causing persistent VAP. This sets the pace for urgent strategies effectively tackling ETT bioburden and curbing the risk of VAP. Preventing microbial colonization has long been recognized as the best approach to avoid biofilm formation and thus, VAP. In the last few decades, technology has advanced, although at a slow pace, regarding the development of new active and/or passive materials for improving ETT surfaces [19]. This relatively slow pace has been shown not to be attributable to a shortage of new ideas, as recently evidenced [19], but to the challenge of overcoming fundamental hurdles associated with VAP (e.g., persistent colonization/biofilm formation; mechanical ventilation time; ICU length of stay; VAP occurrence). In addition, the design and implementation of new biomaterials often imply following specific criteria (e.g., antimicrobial activity; safety-to-use; long-term stability; surface physicochemical properties) to achieve clinical successfulness. This study, which links up with our previous report showing the potential of the immobilization of CHX using a one-step pDA-functionalization approach in the context of orthopedic devices [42], aimed to improve PVC-ETT design by consistently employing dopamine strategy in the immobilization of CHX and CIP on PVC surfaces. A deep characterization regarding surface properties (topography; roughness; wettability), the release profiles, and the potential for reducing microbial colonization while retaining the safety of use was made, so that we can go further with a great ETT coating near achieving success in *vivo*.

The extent of microbial adhesion and subsequent formation of biofilms is highly reliant on certain properties of the bulk polymeric substrates, such as wettability, roughness, and topography [43]. Our prepared coatings imparted PVC surfaces with an altered homogeneously scattered morphology and varied wettability but similar topography (low roughness) to pDA-coated surfaces. However, CIP/CHX co-immobilization yielded coatings with homogenous, hydrophobic, and smooth topographic properties comparable to surfaces immobilized with CHX alone. This result is in accordance with Daud et. al that reported an increase in the hydrophobicity of stainless-steel surfaces after CHX immobilization using a pDA-based approach [44].

In the design of antimicrobial coating strategies, the antimicrobial activity may be addressed either by a direct contact-killing approach, in which the antimicrobial compound is permanently attached to a surface or by a strategy in which the antimicrobial agent is released [45]. Although the first strategy has the advantage of decreasing the propensity for cytotoxicity and development of microbial resistance, there is a major setback associated with its application, namely the fact that the first adherent dead cells may serve as a platform for the adhesion of the next ones, masking the antimicrobial effect of the surface.

This is of particular concern if we bear in mind that the ETT biofilms, which often lead to severe VAP, are comprised of many different phylogenetic organisms and develop quickly after intubation [35,46]. We, therefore, hypothesize that a dual-drug release appears to hold promise, in that it would benefit from a multitude of features, such as a long-lasting and broad-spectrum activity, thereby declining the emergence of antimicrobial resistance, reduction in antimicrobial doses, and good biocompatibility [47]. Our findings showed a weighted but sustained release of CIP and CHX when immobilized on PVC surfaces, with both compounds (alone and combined) retaining contact-killing activity for a long-term period (up to 10 days). Unlike a variety of emergent ETT antimicrobial coatings that have not confirmed long-term stability, our dual-drug coating drives potential application for longer than 24 h, mitigating device failure for longer time periods and, consequently, reducing the real risk for recurrent ETT exchanges and the propensity for developing VAP.

Biocompatibility, as well as stability, is another feature often dismissed in reports addressing antimicrobial materials for ETTs [19]. Despite the efforts that have been made to study both of these traits, data remain scant or controversial, thereby leading to inconclusive safety and clinical usefulness of most emergent ETT coatings. It is the case, for instance, of silver-based ETT coatings. Even though they have rapidly reached the market with promising antimicrobial effectiveness [48,49], there is increasing evidence of the toxicity and short stability attained by released silver ions, in addition to the health concerns regarding the promotion of resistance in clinical isolates [50,51]. When designing a dual-drug coating, in particular, it is imperative that the release of compounds will be assured in doses higher enough to provide an antimicrobial effect but not so that it will be cytotoxic. In this study, coatings incorporating only CIP or CHX did not exhibit toxicity toward mammalian cells, even for the highest doses tested (2 mg/mL). In turn, surfaces co-immobilizing CIP/CHX caused A549 toxicity in a dose-dependent manner, and biocompatibility was only afforded for coatings incorporating antimicrobials with doses up to 0.5 mg/mL. Based on these findings, further antibiofilm studies were performed for coatings displaying no toxic effects against lung epithelial cells.

A key but considerably often-neglected factor has been the ETT biofilm and its polymicrobial etiology. Most antimicrobial coating strategies reported to fight VAP are tested against microorganisms individually [52,53]. To the best of our knowledge, a sole polyurethane-based hydrogel ETT coating incorporating a lead ceragenin (CSA-131) was found to address performance against polymicrobial, including bacterial–bacterial (*P. aeruginosa*/MRSA) and bacterial-fungal (*P. aeruginosa*/*Candida auris*) biofilms [54]. In addition, this latest report reported less efficient antimicrobial performance against polymicrobial biofilms compared to single-species consortia [55]. The present study represents a step forward towards a collective representation of VAP pathogens, while devising an antimicrobial coating strategy to endow the surface of ETTs with the ability to resist the colonization of Gram-negative (*P. aeruginosa*, *K. pneumoniae*, and *A. baumannii*) and Gram-positive bacteria (*S. aureus*, *S. epidermidis*) but also fungi (*C. albicans*), in single, dual and triple-species consortia, thus reflecting worst-case scenarios in the context of VAP. PVC coatings with CHX immobilized alone proved not to be effective against important Gram-negative VAP-relevant bacteria, namely *P. aeruginosa* and *A. baumannii*. In previous studies, ETTs impregnating CHX, particularly combined with other antiseptic agents, had demonstrated potential in reducing colonization by MDR gram-negative bacteria but also by Gram-positive and *Candida* spp. [24,25]. The use of another antimicrobial compound, CIP, showed auspicious in broadening the antimicrobial spectrum of our coating strategy. CIP affords antibacterial effect by binding to bacterial enzymes, DNA gyrase and topoisomerase IV, resulting in permanent double-stranded DNA and cell death [56]. Co-immobilization of CIP and CHX (CIP/CHX) imparted PVC surfaces with the ability to effectively reduce or prevent microbial colonization and biofilm formation by all five bacterial species investigated (*P. aeruginosa*, *A. baumannii*, *K. penumoniae*, *S. aureus*, *S. epidermidis*). In addition to increasing the spectrum of action, reducing antimicrobial resistance and toxicity, a combination of antimicrobials comprises additional advantages, that include rejuvenation of old antibiotics or drug synergism [47] . In fact, CIP/CHX-modified surfaces exhibited both synergic and additive effects against all species, with reduced concentrations of both compounds crucial to assure no toxicity against lung epithelial cells.

The proposed coating strategy was, therefore, further evaluated against polymicrobial consortia, namely involving two and three species. Promising led to conclude that CIP/CHX mixed coating strategy exhibited similar or even better antimicrobial effects against polymicrobial communities when compared to their single-species colonization. The microbiome of VAP has been reported to be mainly comprised of bacteria and only a small percentage of viruses and fungi [57]. The role played by such lesser common species, namely fungal species such as *C. albicans*, on the adverse clinical outcomes in patients with VAP has not been fully elucidated. First, it was assumed that the presence of *Candida* spp. in respiratory tract specimens should be considered colonization rather than infection [58]. In fact, ESCMID guidelines for the management of *Candida* spp. do not recommend antifungal therapy unless there is clear histological evidence of infection [59]. However, other studies have suggested an association between *Candida* spp. colonization and longer mechanical ventilation [60], increased risk for multi-drug resistant bacteria isolation [61], and death [62]. In the present study, *C. albicans* colonization as single species was not impaired as efficiently as the biofilms of bacterial species. However, its presence did not compromise the antibiofilm activity against two major VAP pathogens, *P. aeruginosa* or *S. aureus*, when the coating strategies were evaluated against polymicrobial consortia.

## 5. Conclusions

In this study, the combination of CIP and CHX, using dopamine chemistry, resulted in a dual-drug coating with antimicrobial activity towards a wide spectrum of microorganisms commonly associated with VAP without compromising the viability of lung epithelial cells. Furthermore, a safe-by-design approach was followed, as evidenced by the reduction of the concentration of both compounds so that a balance between antimicrobial features and biocompatibility could be obtained. Worthy of note, the sustained antimicrobial release for a period of up to 10 days strongly evidences the long-term stability and robustness of the antimicrobial immobilization in our dual-drug coating.

Taking all findings together, our dual-drug coating approach holds great potential to be further applied in the next generation of EETs, with promise laying in reducing the incidence rates of VAP.

## ASSOCIATED CONTENT

### Supporting Information

Table S1. Values of minimum inhibitory concentration (MIC) and minimum microbiocidal concentration (MMC) for chlorhexidine (CHX) and ciprofloxacin (CIP), expressed in mg/L, against planktonic cultures of P. aeruginosa, A. baumannii, K. pneumoniae, S. aureus, S. epidermidis, and C. albicans. The antimicrobial susceptibility of planktonic cultures was determined through the broth microdilution method, following the standard European Committee on Antimicrobial Susceptibility Testing (EUCAST) guidelines. (PDF)

Table S2. Determination of possible occurrence of facilitation or synergism for co-immobilization of CIP and CHX at a concentration of 0.5 mg/mL for single-species biofilms formation. The combinations where facilitation or synergism outcomes were obtained are highlighted in bold. In these equations, C refers to the microbial density obtained in the control (PVC surfaces) and S_CHX_, S_CIP_ and S_MIX_ to the surviving cell density after being in contact with surfaces functionalized with CHX, CIP, and the combination of CHX and CIP (MIX). (PDF)

Table S3. Determination of possible occurrence of facilitation or synergism for co-immobilization of CIP and CHX at a concentration of 0.5 mg/mL for dual-species biofilms formation. The combinations where facilitation or synergism outcomes were obtained are highlighted in bold. In these equations, C refers to the microbial density obtained in the control (PVC surfaces) and S_CHX_, S_CIP_ and S_MIX_ to the surviving cell density after being in contact with surfaces functionalized with CHX, CIP and the combination of CHX and CIP (MIX). *(PDF)*

Figure S1. Surface roughness. AFM images of PVC surfaces before and after different pDA-based coating strategies for the immobilization of CIP and/or CHX: (A) unmodified PVC surface; (B) pDA coating; (C) CHX-modified surface with CHX at 0.5 mg/mL; (D) CHX-modified surface with CHX at 2 mg/mL; (E) CIP-modified surface with CIP at 0.5 mg/mL; (F) CIP-modified surface with CIP at 0.5 mg/mL; (G) CIP/CHX-modified surface with CIP and CHX at 0.5 mg/mL; (H) CIP/CHX-modified surface with CIP and CHX at 2 mg/mL. (PDF)

Figure S2. Effect of different selective media on bacterial and fungal growth. A microbial suspension of each microorganism investigated was adjusted to 1x10^6^ CFU/mL and plated on the different selective media. (PDF)

## AUTHOR INFORMATION

### Author Contributions

The manuscript was written through contributions of all authors. All authors have given approval to the final version of the manuscript.

### Notes

The authors declare no competing financial interest.

## ACKNOWLEDGMENTS

This work was supported by the Portuguese Foundation for Science and Technology (FCT) under the scope of the strategic funding of UIDB/04469/2020 unit, and by LABBELS – Associate Laboratory in Biotechnology, Bioengineering and Microelectromechanical Systems, LA/P/0029/2020. The authors also acknowledge the support, through the Programa Operacional Competitividade e Internacionalização (COMPETE2020) and by national funds, through the FCT, of the POLY-PrevEnTT project (PTDC/BTMSAL/29841/2017-POCI-01-0145-FEDER-029841). A special thanks to Professor Alexandr Nemec from National Institute of Public Health in Prague, for kindly providing the *A. baumannii* strain used in this study.

