## Supplemental material for "Co-immobilization of ciprofloxacin and chlorhexidine as a long-term, broad-spectrum antimicrobial dual-drug coating for polyvinyl chloride (PVC)-based endotracheal tubes"

[1] Current affiliation: INL - International Iberian Nanotechnology Laboratory, Av. Mestre José Veiga, 4715-330 Braga, Portugal

**Table S1**. Values of minimum inhibitory concentration (MIC) and minimum microbiocidal concentration (MMC) for chlorhexidine (CHX) and ciprofloxacin (CIP), expressed in mg/L, against planktonic cultures of *P. aeruginosa*, *A. baumannii*, *K. pneumoniae*, *S. aureus*, *S. epidermidis*, and *C. albicans*. The antimicrobial susceptibility of planktonic cultures was determined through the broth microdilution method, following the standard European Committee on Antimicrobial Susceptibility Testing (EUCAST) guidelines.

| | **CHX** | | **CIP** | |
|---|---|---|---|---|
| | **MIC** | **MMC** | **MIC** | **MMC** |
| ***P. aeruginosa*** | 3.13 - 6.25 | 12.5 - 25 | 0.25 | 0.5 - 1 |
| ***A. baumannii*** | 12.5 - 25 | 25 | 2 | 4 - 8 |
| ***K. pneumoniae*** | 0.78 - 1.56 | 3.125 | <0.016 | <0.016 |
| ***S. aureus*** | 0.78 - 1.56 | 1.56 - 6.25 | 0.5 | 0.5 - 1 |
| ***S. epidermidis*** | 0.78 - 1.56 | 1.56 - 6.25 | 0.25 | 0.5 - 1 |
| ***C. albicans*** | 6.25 - 12.5 | 6.25 - 25 | >64 | >64 |

**Table S2. Determination of possible occurrence of facilitation or synergism for co-immobilization of CIP and CHX at a concentration of 0.5 mg/mL for single-species biofilms formation.** The combinations where facilitation or synergism outcomes were obtained are highlighted in bold. In these equations, C refers to the microbial density obtained in the control (PVC surfaces) and $S_{CHX}$, $S_{CIP}$ and $S_{MIX}$ to the surviving cell density after being in contact with surfaces functionalized with CHX, CIP, and the combination of CHX and CIP (MIX).

| **Microorganism** | **Synergism** [Log ($S_C$)– Log ($S_{CHX}$) – Log ($S_{CIP}$) + Log ($S_{MIX}$)] | **Facilitation** [Log ($S_{MIX}$)– Log ($S_{CHX}$) / Log ($S_{MIX}$) - Log ($S_{CIP}$)] |
|---|---|---|
| *P. aeruginosa* | **-4.908** | **-6.144 / -5.233** |
| *A. baumannii* | **-1.933** | **-4.161 / -1.885** |
| *K. pneumoniae* | 5.393 | **-0.167 / -0.083** |
| *S. aureus* | 1.859 | **-2.514 / -1.962** |
| *S. epidermidis* | 6.534 | **-0.620 / -0.033** |
| *C. albicans* | **-0.557** | **-0.283 / -0.753** |

**Table S3**. **Determination of possible occurrence of facilitation or synergism for co-immobilization of CIP and CHX at a concentration of 0.5 mg/mL for dual-species biofilms formation** The combinations where facilitation or synergism outcomes were obtained are highlighted in bold. In these equations, C refers to the microbial density obtained in the control (PVC surfaces) and $S_{CHX}$, $S_{CIP}$ and $S_{MIX}$ to the surviving cell density after being in contact with surfaces functionalized with CHX, CIP and the combination of CHX and CIP (MIX).

| Dual-species consortia | Synergism<br>[Log ($S_C$)– Log ($S_{CHX}$) – Log ($S_{CIP}$) + Log ($S_{MIX}$)] | Facilitation<br>[Log ($S_{MIX}$)– Log ($S_{CHX}$) / Log ($S_{MIX}$) - Log ($S_{CIP}$)] |
|---|---|---|
| *P. aeruginosa* | **-1.797** | **-6.449 / -2.040** |
| *K. pneumoniae* | 4.732 | **0 / 0** |
| *P. aeruginosa* | **-3.677** | **-7.349 / -4.138** |
| *S. aureus* | 0.366 | **-3.277 / -2.679** |
| *P. aeruginosa* | **-3.544** | **-6.791 / -3.797** |
| *S. epidermidis* | 4.950 | **-1.109 / 0.094** |
| *P. aeruginosa* | -2.909 | **-7.267 / -2.579** |
| *C. albicans* | -0.853 | **-1.853 / -1.711** |

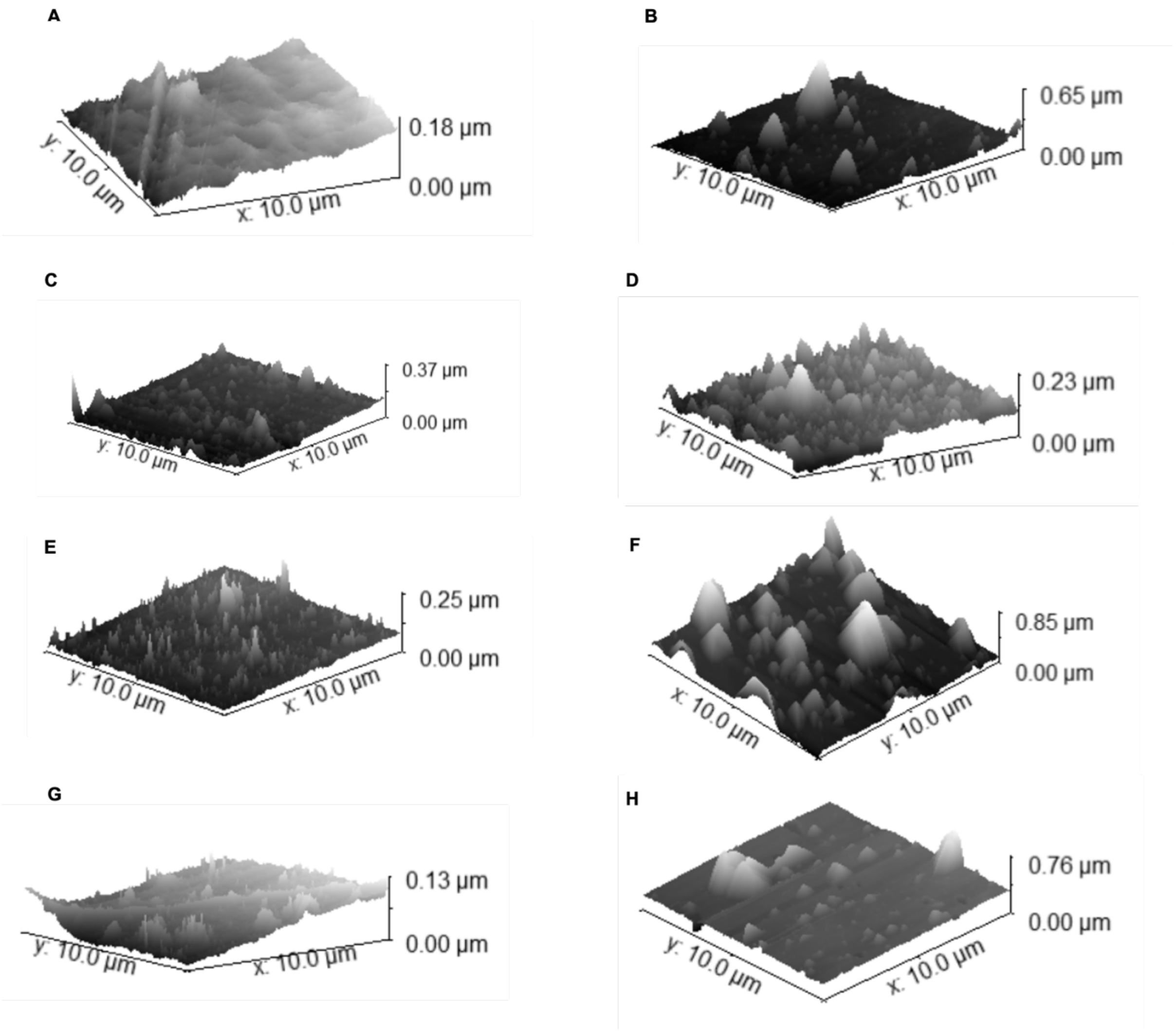


**Figure S1. Surface roughness.** AFM images of PVC surfaces before and after different pDA-based coating strategies for the immobilization of CIP and/or CHX: (A) unmodified PVC surface; (B) pDA coating; (C) CHX-modified surface with CHX at 0.5 mg/mL; (D) CHX-modified surface with CHX at 2 mg/mL; (E) CIP-modified surface with CIP at 0.5 mg/mL; (F) CIP-modified surface with CIP at 0.5 mg/mL; (G) CIP/CHX-modified surface with CIP and CHX at 0.5 mg/mL; (H) CIP/CHX-modified surface with CIP and CHX at 2 mg/mL.

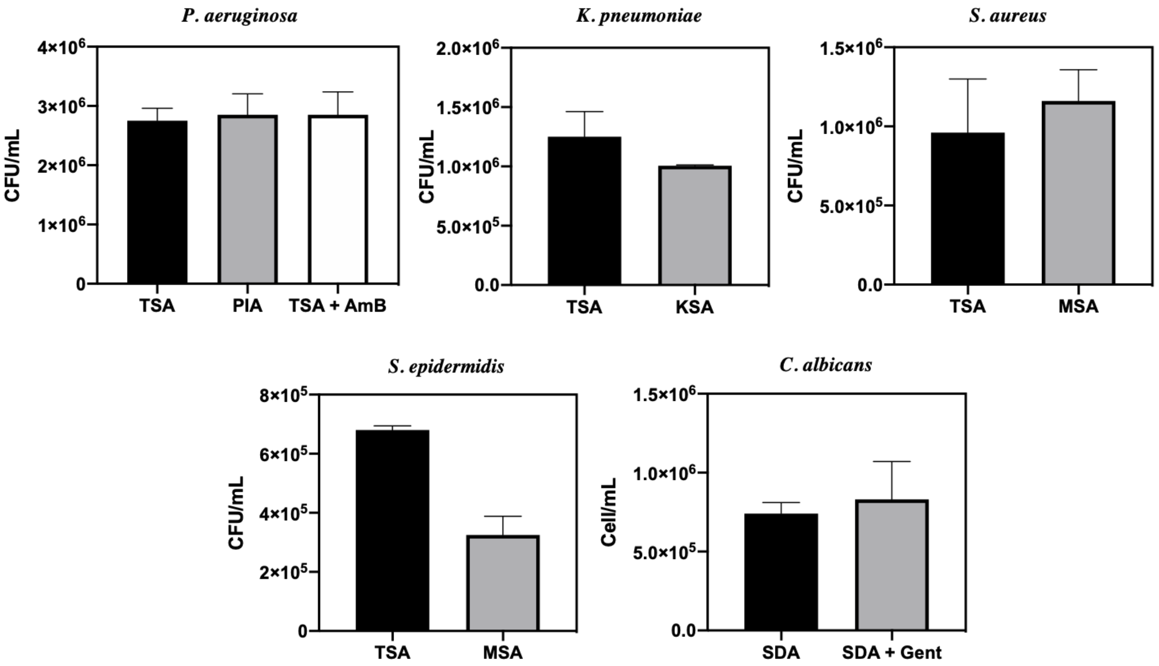


**Figure S2. Effect of different selective media on bacterial and fungal growth.** A microbial suspension of each microorganism investigated was adjusted to $1x10^6$ CFU/mL and plated on the different selective media.
